# Model-Based Estimation of Active and Passive Muscle Forces Using MRE in Forearm Muscles During 2-DOF Wrist Tasks

**DOI:** 10.1101/2024.02.15.580561

**Authors:** Cody A Helm, Fabrizio Sergi

**Affiliations:** Human Robotics Laboratory, Departments of Biomedical and Mechanical Engineering, University of Delaware, Newark, DE, 19713

**Author notes:** Corresponding author: Fabrizio Sergi.

## Abstract

Magnetic resonance elastography (MRE)-based muscle force estimation methods have been proposed to estimate individual muscle forces based on measurements of shear wave speed and joint torque in multiple postures. To estimate both the slope and offset parameters of the relationship between shear wave speed and muscle force, it is necessary to collect measurements in a plethora of postures in case of substantial muscle redundancy. However, anisotropic MRE requires a structural and diffusion-tensor imaging scan in each posture, which is infeasible given the time constraints of MRI imaging.

The objectives of this work were to develop a muscle force estimator with sufficient accuracy that would only require a limited set of postures, and to evaluate its effectiveness under a variety of measurement conditions. We developed a novel MRE-based muscle force estimator, which decouples shear wave speed into its active and passive components and solves for the slope and offset parameters, independently. We assessed the effectiveness of the proposed estimator under different simulated measurement conditions, with varying levels of noise, and compared it to the original MRE-based estimator.

The proposed estimator results in a reduction in the estimation error for the offset parameter, for all muscles, without significant degradation in the estimation error for the slope parameter. However, the proposed muscle force estimator does not improve the goodness-of-fit or the cross-validation error compared to the original estimator. In conclusion, the proposed MRE-based muscle force estimator improves the estimation of muscle-specific parameters and may yield increased muscle force estimation performance.

## I. INTRODUCTION

Knowledge of how the musculoskeletal system distributes joint load amongst individual muscles is one of the grand challenges of biomechanics. Insight of individual muscle force contributions would provide guidance into the design of human-interacting robotic systems and prostheses [1], experimental validation for musculoskeletal models [2], and an improved understanding of motor impairment diagnosis and progression in musculoskeletal injuries [3].

One method to quantify individual muscle force involves muscle activity measurements with muscle force calibration procedures. This method combines surface electromyography (sEMG) recordings from individual muscles with joint torques and a musculoskeletal model to compute estimates of individual muscle force [4]. These methods have shown promise in both static and dynamic muscle contractions [5]. However, in wrist tasks where non-superficial muscles contribute substantially to the resultant joint torque, sEMG-based estimates of muscle force can be significantly inaccurate due to the presence of un-measurable function from non-superficial muscles and cross-talk between adjacent muscles [6]. Ultrasound elastography, a shear wave elastography (SWE) technique, has been developed to quantify mechanical properties of soft tissues. Ultrasound elastography can quantify the mechanical properties of individual muscles [3], [7], and establish the relationship between shear wave speed and skeletal muscle force undergoing static and dynamic contractions. While ultrasound elastography offers excellent temporal resolution, it is also limited in its ability to measure the shear wave speed in non-superficial muscles.

Magnetic resonance elastography (MRE) incorporates MRI imaging to measure the shear wave speeds of skeletal muscles in a 3D tissue volume. This enables to measure the muscle shear stiffness at a larger field-of-view. Shear wave speed of upper extremity muscles such as the biceps brachii [8] and the flexor digitorium profundus (FDP) have been quantified with MRE during rest and active contractions. MRE is sensitive to detect differences in muscle mechanical properties associated with injured muscles compared to healthy muscles [9]. Recently, our group has shown that MRE can estimate individual muscle forces in the forearm in humans during isometric wrist contractions based on repeated measurements in different postures [10]. This work has shown that shear wave speed squared measurements can explain 70% of the variance in wrist torque via a linear regression model and can back estimate wrist torque with 0.3 Nm average torque error [10].

One limitation of most previous muscle MRE work is the assumption of an isotropic material in estimating the relationship between the measured displacement field and the shear wave modulus, *G*. New methods for anisotropic MRE imaging have been proposed [11], and they account for the mechanical properties of muscle fibers when solving the nonlinear inversion algorithm and obtaining the shear stiffness 3D tissue maps. On the one hand, MRE protocols based on a larger number of postures are advantageous because they allow to differentiate the independent contribution of multiple agonist and antagonist muscles acting around the wrist joint to produce joint torque. On the other hand, however, the use of a large number of postures in an MRE protocol may be non-viable experimentally due to the need of collecting anatomical and diffusion tensor imaging (DTI) in multiple postures, leading to participant fatigue. Additionally, current MRE-based muscle force estimators provide acceptable accuracy in estimating the slope coefficient but are inaccurate in estimating the intercept because they assume it to be zero [6]. As such, identification of force estimation algorithms that are accurate in estimating muscle force from MRE data under a limited number of postures would allow to reap the benefits of anisotropic MRE protocols. Additionally, an improved estimation of passive muscles forces may reduce the average wrist torque error observed in previous MRE-based muscle force estimation [10].

Towards the goal of enabling the use of anisotropic MRE protocols for estimating individual muscle force, this paper presents the development of a new muscle force estimator that can be adequately combined with anisotropic MRE to accurately estimate the parameters of the shear-wave speed to force relationship for all thirteen forearm muscles and evaluate its effectiveness in estimating muscle force under a variety of experimental conditions.

## II. Materials and Methods

### A. Muscle Mechanics

Our previous work reviews the derivation of the mathematical relationship between shear wave speed measurements and muscle forces [10]. Here, we will provide a brief summary of the derivation to enable us to use this relationship in the muscle force estimator.

The outcome from MRE data are muscle-specific measurements of shear-wave speed *v*_*SH*_. The relationship between shear-wave speed and muscle force can be derived by combining the relationship between muscle force (*f*_*MT*_) and axial load (*σ*), the relationship between Young’s modulus (*E*) and shear modulus (*G*), and the relationship between shear wave speed, axial load, and shear modulus. Muscle force is related to the axial load in the muscle through its cross-sectional area (*A*_*CS*_).

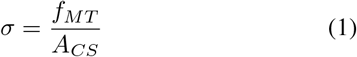

In muscle tissue, *E* increases linearly with the axial stress (*σ*) from a starting unstressed value (*E*_0_) with a slope of *α* [12].

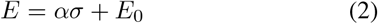

It has been shown that there is a linear relationship between Young’s modulus (*E*) and shear modulus for skeletal muscles [12]. Thus, with knowledge of a linear proportionality constant, *β*, we can obtain the shear modulus, *G*, as

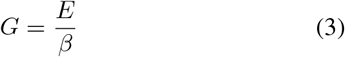

Shear-wave speed is a function of both shear modulus, *G*, and axial load, *σ*, for axially isotropic materials. Martin et al., [7] provides a relationship between shear-wave speed squared 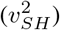, shear modulus, and axial load:

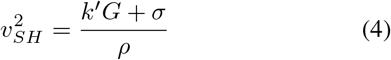

Thereby, substituting (1), (2), and (3) into (4), we obtain a relationship between 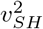 and muscle force. The resulting equation is a linear relationship between muscle force and 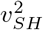 with a slope *γ* and offset *κ*:

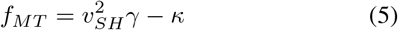

Thus, muscle forces can be obtained from 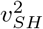 measurements once the muscle-specific parameters *γ* and *κ* have been determined.

### B. Muscle Force Estimation

To determine the parameters *γ* and *κ* for the musclespecific relationship between 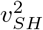 and muscle force, we formulated a muscle force estimation procedure that extends the one presented in previous work [6]. The muscle force estimator combines measurements of 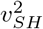, wrist torques, and wrist postures with parameters extracted from a musculoskeletal model [13], which includes *m* muscles spanning the wrist joint. A muscular Jacobian matrix, **J**, was obtained for each of the *m* muscles and considered wrist postures from the musculoskeletal model via OpenSim [6]. Components of matrix **J** are the moment arm values with each row corresponding to one of the *n* wrist DOFs, and each column corresponding to a specific muscle, resulting in **J** being a *n* × *m* matrix. In this work, we consider *n* = 2, as we consider torque production in the flexion/extension (FE), and radial/ulnar deviation (RUD) axes. For estimation, a *n* × 1 wrist torque vector ***τ*** = [*τ*_*F E*_; *τ*_*RUD*_] measured during each isometric contraction using the MREbot [14] is related to the *m* × 1 muscle force vector as:

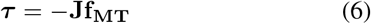

By substituting the relationship between muscle force and shear-wave velocity into the previous equation, we obtain the relationship between the measured 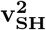 and the measured ***τ*** :

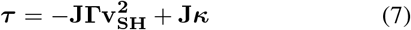

where 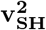 is a *m* × 1 vector that contains the measurements of 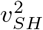for each muscle, **Γ** = diag(*γ*_1_, *γ*_2_, …, *γ*_13_) is a *m* × *m* matrix that contains muscle-specific slope constants in its diagonal elements, and ***κ*** is a *m* × 1 vector that contains the muscle-specific offset constants.

Overall, the equation above is a linear equation of the form ***X***_***l***_***β*** −***y***_***l***_ = 0, where 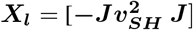 is a *n* × 2*m* matrix obtained by horizontal concatenation of a *n* × *m* matrix with elements 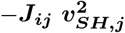 and the ***J*** matrix; ***β*** = [***γ***; ***κ***] is a 2*m* × 1 vector of unknowns obtained by vertical concatenation of the unknown parameter vectors ***γ*** = (*γ*_1_, *γ*_2_, …, *γ*_*m*_)^*T*^ and ***κ*** = (*κ*_1_, *κ*_2_, …, *κ*_*m*_)^*T*^ and ***y***_***l***_ = ***τ***_***l***_, and subscript *l* indicates each contraction. By collecting data during *N* isometric contractions at different wrist postures, it is possible to construct a *nN* × 2*m* matrix ***X*** = [***X***_**1**_; ***X***_**2**_; …; ***X***_***N***_] via vertical concatenation of the contraction-specific ***X***_***l***_ matrices to account for the *nN* measurements ***y*** = (***τ***_1_, ***τ***_2_, …, ***τ***_*l*_)^*T*^.

Using least squares regression, it is then possible to estimate the vectors 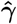 and 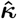 given measurements of joint torque, 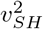, and knowledge of the moment arm matrix ***J*** from a musculoskeletal model. After estimating 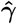 and 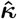, the estimated parameters can be used to convert measurements of 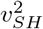 to estimates of muscle force using (5).

The second term ***J*** in the matrix ***X***_***l***_ is a *n* × *m* matrix (in general, a wide matrix: *m > n* due to muscle redundancy) whose elements are obtained for each wrist posture. The same ***J*** is used for all contractions that occur in a single posture. Therefore, the number of linearly independent rows in this matrix is a maximum of *n*, or 2. For an experimental protocol with *P* postures, the rank *r* of the experimental matrix **X**, is defined as:

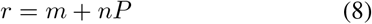

The *r* also corresponds to the number of non-zero parameters that can be estimated in the least-square sense. For all parameters in ***β*** to be estimated, *r* ≥ 2*m*.

For the case of forearm MRE, *m* = 13 and *n* = 2, so this algorithm would require to collect data in seven wrist postures for the experimental matrix to be full-rank, i.e. *r >* 2*m*. For anisotropic MRE muscle imaging protocols, structural MRI images and a DTI scan are required to define muscle segments and tract the muscle fibers in each additional wrist posture. Due to the time-constraints of MRI experiments, it is not possible to collect MRE measurements in seven different wrist postures in a single session without inducing significant effects of participant fatigue, or requiring structural images to be collected in separate sessions. Therefore, there is a need to develop a muscle force estimator to allow for estimating all unknown parameters even in the presence of a limited number of wrist postures.

### C. Proposed Muscle Force Estimator

The proposed muscle force estimator aims to estimate both 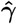 and 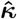 for all muscles by decoupling equation (7) into its active and passive components to solve for each parameter individually. First, the set of 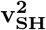 measurements is decoupled into active and passive components. The passive 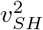 matrix 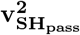 is computed for each posture as a *P* × *m* matrix. Each row *p* of 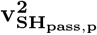 is computed as the average of 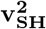 values measured over multiple repetitions *k* during rest for each posture *p*:

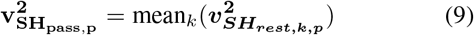

The active 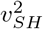 matrix, 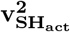, is *N* × *m*, where *N* is the number of contractions. The rows of 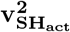 are obtained by subtracting the posture-specific 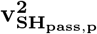 from the measured 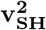 in that posture for all *N* contractions. For each posture *p* and contraction *l*,

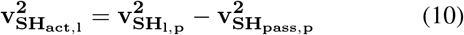

Wrist torques ***τ*** are also decoupled into active and passive components. For each contraction, the passive joint torque, ***τ***_***pass***_, is a *n* × 1 vector resulting from passive muscle forces, **f**_**pass**_, which are posture-specific and estimated from a musculoskeletal model [13].

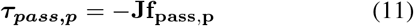

During MRE data collections, at each new posture the force/torque sensor is zeroed with the participant at rest to remove any non-zero torque values resulting from the contribution of passive muscle force in that posture [10]. Thus, the active torque, ***τ***_***act***_, is a *n* × 1 vector equal to the measured torque values, ***τ***_***m***_. Parameter estimation is performed using first the active torque values for all *N* contractions to obtain the muscle specific 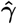 slopes, i.e., by solving the equation below for **Γ** using ordinary least squares regression.

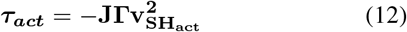

The experimental matrix for this design is a tall matrix with dimensions (*nN*) × *m*, with linearly independent columns, and thus an *m*-rank matrix. As such, it is possible to solve for the muscle-specific slope coefficients for each of the *m* muscles using ordinary least squares.

Once the muscle-specific slope coefficients are determined as 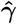, and 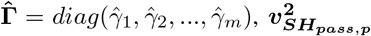 and ***f***_***pass***,***p***_ are used to obtain the 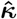 coefficients for all muscles and each posture *p*.

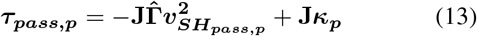

From here, we can substitute (11) in the expression for torque, and then simplify **J** from the equation, solving for the unknown set of parameters 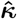.

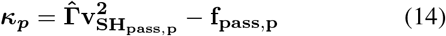

The muscle-specific 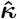 offset terms are determined by taking the average over all wrist postures.

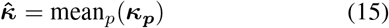

As a recap, the rank of the experimental matrix for the proposed estimator is independent of *P*, thus a protocol with *P* = 1 is sufficient to obtain a full-rank experimental matrix.

### D. Model-Based Simulations of Shear Wave Speed Squared Measurements

We simulated virtual experiments where the quantities of 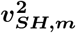 and ***τ***_***m***_ were measured for a given experimental protocol composed of *N* isometric contractions to investigate the accuracy of our proposed estimator compared to the original estimator. Each contraction was defined as an isometric wrist torque applied at a specific wrist posture. For each contraction, we obtained simulated values of muscle forces using an optimization-based redundancy solver that aimed to minimize the global activation level (*GAL* = ∑_*j*_ (*activations*_*j*_)^2^) for a given ***J*** and ***τ***_***m***_. For all contractions, the active simulated force values were added to the posture-dependent passive force extracted via a musculoskeletal model [13]. With known values of 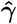 and 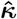, simulated muscle forces were converted to simulated 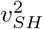 values, and different sources of noise added to the simulated measurement (see below for details). The simulated 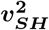 matrix was included with the ***τ***_***m***_ and ***J*** in an ordinary leastsquares regression to obtain muscle-specific estimates of 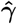 and 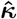.

### E. Evaluation of Estimators

We conducted *N* = 50 virtual experiments where measurements of 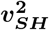 were simulated under different noise levels and experimental protocols defined as a set of wrist torques and wrist postures to evaluate the effectiveness of each estimator. Similarly to our previous work [6], we applied two forms of noise/variance into the simulated 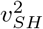 values: 1) Physiological variability, *ϵ*^*A*^, was implemented to account for non-optimal muscle contraction patterns that can be used to achieve the desired wrist torque; 2) Measurement noise, *ϵ*^*M*^, was implemented to account for variance in data quality. For each virtual experiment, *ϵ*^*M*^ and *ϵ*^*A*^ were selected from uniform distributions with *ϵ*_*max*_ ∈ [0 0.5 1], i.e., adding error up to 100% of the true value of force. The distribution for 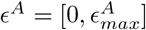 and 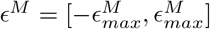. We considered two models, the original muscle force estimator and the proposed muscle force estimator. For all virtual experiments, we included the measurements from all thirteen forearm muscles to reproduce conditions of MRE imaging.

We considered three different experimental protocols defined based on wrist postures and repetitions. We simulated virtual experiments for a five posture / one repetition (5P/1R), a two posture / one repetition (2P/1R), and a two posture / three repetition (2P/3R) protocol. The wrist postures for the five posture protocol were defined as ([*θ*_*F E*_, *θ*_*RUD*_] ∈ [−30, 0], [−15, 0], [0, 0], [15, 0], [30, 0]^°^) and the wrist postures for the two posture protocol were defined as [− 15, 0], [15, 0]^°^. Desired wrist torques were defined as unitary magnitude in all four cardinal directions, flexion, extension, radial deviation and ulnar deviation, and a rest condition. For all contractions, we simulated 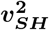 measurements with *γ* = 10 and *κ* = 500 for all muscles.

For the muscle level analysis, we obtained estimates of 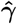 and 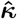 using both the original and proposed estimator in all protocols and noise combinations. We quantified the accuracy of each estimator by computing the percent error in estimating 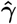 and 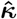 for all *m* forearm muscles. 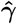 and 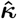 error for the five main wrist muscles (extensor carpi radialis longus and brevis (ECRL, ECRB), extensor carpi ulnaris (ECU), flexor carpi radialis (FCR), and flexor carpi ulnaris (FCU)) were computed by taking the median percent error over those muscles.

Under the model assumptions, the force measured from multiple muscles spanning the wrist joint should sum as described in (6) to yield the measured value of joint torque accounting for passive joint torque. In other words, if 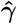 and 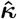 are the estimated muscle-specific parameters and 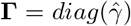, the joint torques resulting from the estimated muscle forces 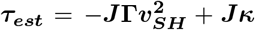 should be equal to joint torque values ***τ***_***m***_ measured via the F/T sensor along with the passive wrist torques. To quantify how well the estimators can reconstruct the joint-level torque, we obtained *R*^2^ goodness of fit values for each virtual experiment and estimator type. For both estimators, the *R*^2^ value was obtained by comparing the back-estimated torques predicted by each estimator, from eq. 7, and the desired torques.

Moreover, since the estimators use a different set of parameters, and are thus not directly comparable based on *R*^2^, we also performed a leave-one-out cross validation (LOOCV) analysis. We quantified the LOOCV error as the unsigned difference between the estimated torque value of the unseen wrist contraction and the desired wrist torque for that contraction. For each virtual experiment, we computed the average LOOCV error as the average over all *N* contractions. We performed a two-way rmANOVA with fixed factors of estimator and noise level, grouped by protocol, with outcomes *R*^2^ and LOOCV error. Posthoc Tukey HSD paired-comparisons was performed with a significance threshold of *α* = 0.05. Then, as a proxy to quantify the accuracy of muscle force estimation, we quantified the torque estimation bias ***b***_***τ***_, separately, for each contraction,

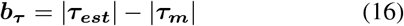

Muscle and joint level analysis results for the original and proposed muscle force estimators for all experimental conditions are shown below.

## A. Muscle Level Analysis

For the wrist muscles, in the highest noise level, the original estimator yields 0.18, 0.44, and 0.19 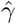 error for the 5P/1R, 2P/1R, and 2P/3R protocols, respectively, while the proposed estimator yields 0.18, 0.43, and 0.19 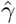 error, respectively (Fig.1). For both estimators, 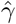 error for wrist muscles increases with *ϵ*^*M*^ but not *ϵ*^*A*^. Additionally, 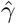 error decreases with an increase in postures and repetitions.

**Fig. 1.**
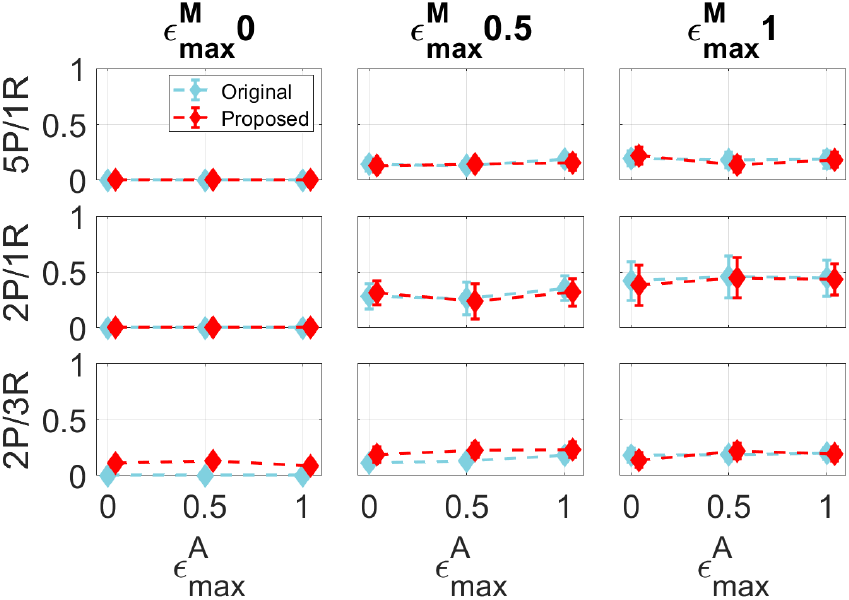
Median 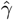 error [n.u.] for wrist muscles grouped by noise condition (columns), protocol (rows), physiological noise (*x* axis), and estimator (overlay). Diamonds indicate the between-runs median and the whiskers extend to *±* 1 standard deviation.

For the wrist muscles, for the highest noise level, the original estimator yields 1.0, 1.0, and 1.0 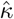 error for the 5P/1R, 2P/1R, and 2P/3R protocols, respectively, while the proposed estimator yields 0.18, 0.47, and 0.23 median 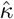 error, respectively, see Fig.2. 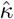 error increases with *ϵ*^*M*^ but not *ϵ*^*A*^ for both estimators. The original estimator results in 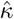 error centered around 1.0, while the proposed estimator results in a lower 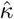 error for all noise levels.

**Fig. 2.**
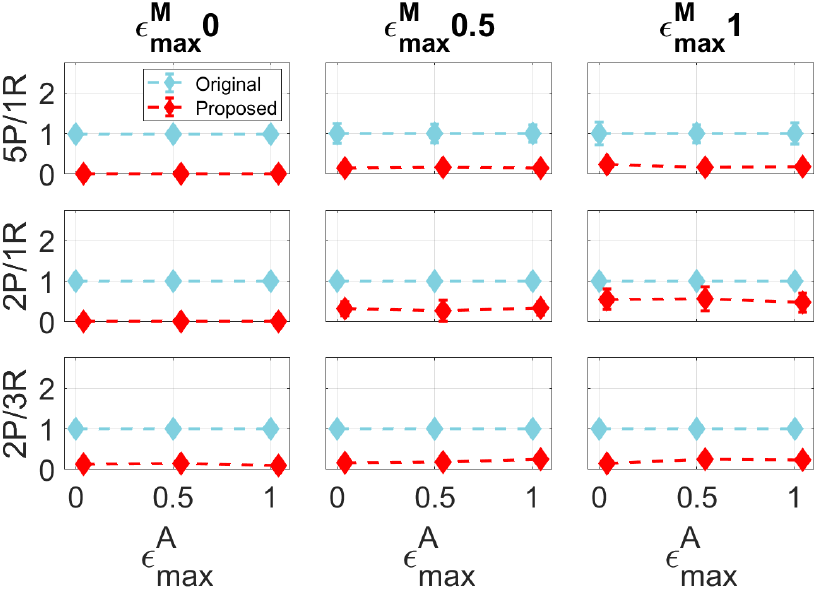
Median 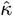 error [n.u.] for wrist muscles grouped by noise condition, protocol, physiological noise, and estimator. Diamonds indicate the between-runs median and the whiskers extend to *±* 1 standard deviation.

### B. Joint Level Analysis

For the highest noise level, the original estimator yields an *R*^2^ of 0.96, 0.99, and 0.95 for the 5P/1R, 2P/1R, and 2P/3R, respectively, see Fig.3. The proposed estimator yields an *R*^2^ of 0.93, 0.90, and 0.93 for the 5P/1R, 2P/1R, and 2P/3R, respectively. *R*^2^ decreases with an increase in *ϵ*^*M*^ for both estimators but does not vary with *ϵ*^*A*^ for either estimator. Statistical analysis showed that the original estimator yielded a significantly higher *R*^2^ value compared to the proposed estimator for high noise levels for the 5P/1R (*t* = 14.56, *p <* 0.001), 2P/1R (*t* = 9.22, *p <* 0.001), and 2P/3R (*t* = 11.82, *p <* 0.001) protocol.

**Fig. 3.**
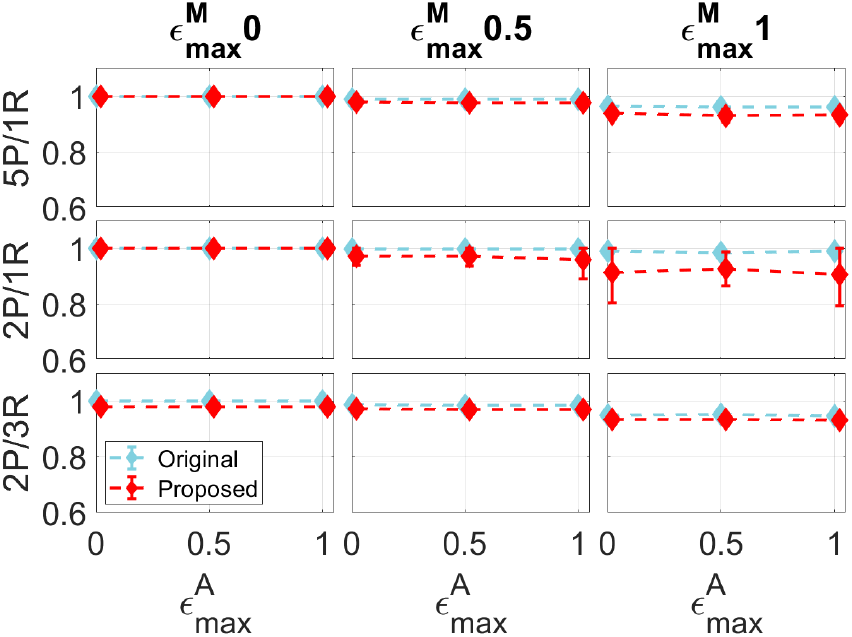
Mean goodness of fit (*R*^2^) values grouped by noise condition, protocol, physiological noise, and estimator. Diamonds indicate the betweenruns mean and the whiskers extend to *±* 1 standard deviation.

For the highest noise level, the original estimator yields an average LOOCV error of 0.30, 1.07, and 0.26 Nm for the 5P/1R, 2P/1R, and 2P/3R, respectively, shown in Fig.4, while the proposed estimator yields an average LOOCV error of 0.27, 0.90, and 0.26 Nm, respectively. Paired comparisons indicate that LOOCV error was not found to be significantly different between the original and the proposed estimator (*t* = 1.26, *p* = 0.207) for high noise conditions with data from all protocols. LOOCV error increases with *ϵ*^*M*^ but does not vary with *ϵ*^*A*^ for both estimators. LOOCV error decreases with an increase of postures and repetitions. For a high noise level, the proposed estimator yields a higher back-estimated torque variation compared to the original estimator for all protocols, see Fig.5. Both FE ***b***_***τ***_ and RUD ***b***_***τ***_ are slightly higher in the proposed estimator than the original in all protocols (Fig.6).

**Fig. 4.**
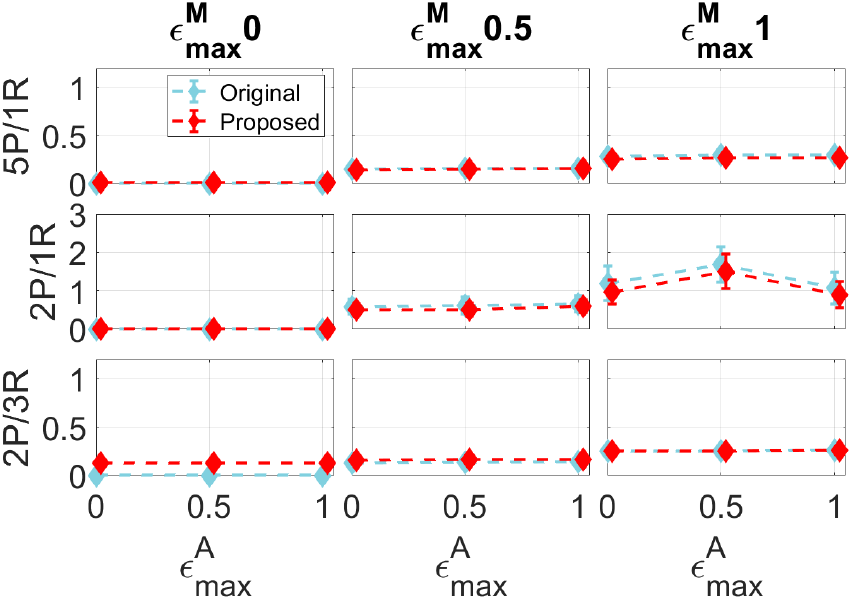
Mean LOOCV error [Nm] values grouped by noise condition, protocol, physiological noise, and estimator. Diamonds indicate the between-runs mean and the whiskers extend to *±* 1 standard deviation.

**Fig. 5.**
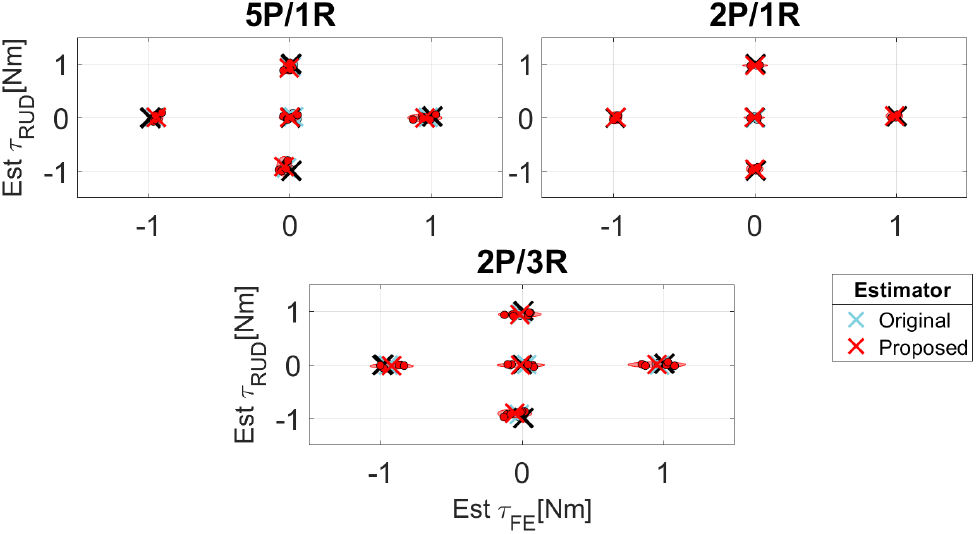
Back-estimated wrist torque mean and 95% confidence intervals plotted for each estimator (overlay) for a high noise level, and protocol (subplots). Black crosses indicate the cued torque value. Each dot represents the between-run average of the back-estimated torque corresponding to a given contraction state.

**Fig. 6.**
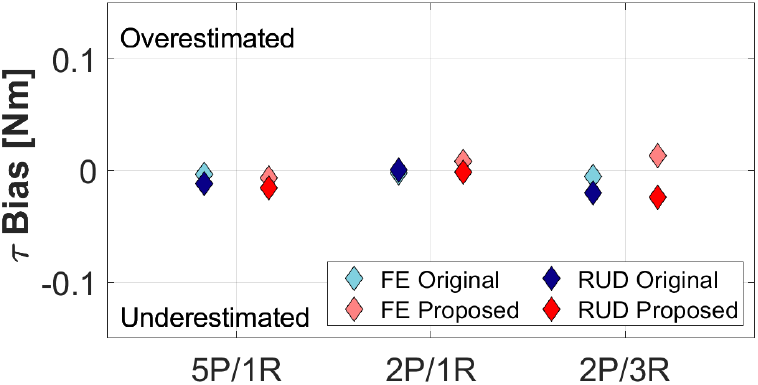
Average torque bias (***b***_***τ***_) [Nm] for each experimental protocol (*x* axis) and estimator (colors), for a high noise level. Negative bias corresponds to an under-estimated wrist torque and positive bias corresponds to an overestimated wrist torque.

## IV. Discussion

The work in this paper presents a novel muscle force estimator for MRE-based measurements that decouples the measured torques and shear-wave speeds to estimate the slope and offset coefficients for the linear relationship between 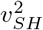 and muscle force. In the proposed estimator, ordinary least squares regression is performed with the measured 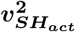 to obtain the slope coefficients for all muscles. Then, the estimated slope coefficients are combined with the measured 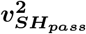 and model-estimated forces to obtain the offset coefficients. We hypothesized that the proposed estimator would improve the accuracy of the muscle force estimation compared to the original estimator. To test this hypothesis, we quantified the estimated parameter percent error for wrist muscles for both estimators. Additionally, we computed the average *R*^2^, average LOOCV error, and the average back-estimated torque bias to evaluate the accuracy of each estimator using analyses that can be replicated experimentally.

Overall, the muscle level analysis shows that the proposed estimator decreases the 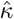 estimation error compared to the original estimator for all protocols at high noise levels while achieving comparable 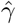 error in the two estimators. 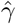 and 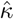 error increase with *ϵ*^*M*^ but not *ϵ*^*A*^. For joint level analysis, the original estimator results in a better *R*^2^ value compared to the proposed estimator. The two estimators yield similar average LOOCV error for high noise combinations. Both *R*^2^ and LOOCV error worsen with an increase in *ϵ*^*M*^ but not *ϵ*^*A*^. The proposed estimator results in a slight increase in ***b***_***τ***_ compared to the original estimator. Taken together, the proposed muscle force estimator improves the overall parameter estimation accuracy, but may reduce the back-estimated torque accuracy compared to the original estimator.

The proposed estimator greatly reduces 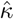 estimation error with a comparable 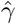 error for the two estimators. As 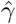 is torque dependent, the capability of estimating 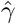 is not and expected to change with decoupling the experimental matrix which is reflected in our findings. For the proposed estimator, 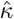 estimation accuracy is dependent on accurate 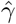 estimation 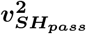. Previous studies have shown accuracy in using rest contractions to estimate the passive muscle force components [10], [15]. The original estimator yields a higher *R*^2^ value than the proposed estimator. One explanation for this is that the original estimator encompasses more parameters when solving the least-squares regression model and therefore can better fit the data to the wrist torques. This is beneficial for *R*^2^ but in cases of high measurement noise can lead to high parameter estimation error. For the LOOCV analysis, the two estimators yield a low average LOOCV error for the 5P/1R and 2P/3R protocols, suggesting that they generalize well to an independent data set.

Back-estimated torque analysis shows that the proposed estimator results in a slight increase in torque bias and variability compared to the original estimator. One explanation might be that in the proposed estimator 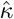 is obtained for each muscle without minimizing the estimated torque residuals. For high noise combinations, the 2P/3R protocol is advantageous in minimizing parameter estimation error and LOOCV error. The proposed estimator offers an improvement in estimating the offset coefficient which can be applied to better quantify passive muscle forces. This work is limited in that the both estimators were only compared using simulated MRE measurements. It is not fully known how the accuracy of each estimator would extend to experimental data. Thus, future directions consists of implementing both estimators to experimental MRE data.

